# Metaproteomic analysis of nasopharyngeal swab samples to identify microbial peptides and potential co-infection status in COVID-19 patients

**DOI:** 10.1101/2023.01.31.525328

**Authors:** Surbhi Bihani, Aryan Gupta, Subina Mehta, Andrew Rajczewski, James Johnson, Dhanush Borishetty, Timothy J. Griffin, Sanjeeva Srivastava, Pratik Jagtap

## Abstract

Respiratory infections disrupt the microbiota in the upper respiratory tract (URT), putting patients at a risk for subsequent infections. During the pandemic, cases of COVID-19 were aggravated by secondary infections because of impaired immunity and medical interventions, which was clearly evident in the second wave of COVID-19 in India. The potential dangers and clinical difficulties of bacterial and fungal secondary infections in COVID-19 patients necessitate microbial exploration of the URT. In this regard, mass spectrometry (MS)-based proteome data of nasopharyngeal swab samples from COVID-19 patients was used to investigate the metaproteome. The MS datasets were searched against a comprehensive protein sequence database of common URT pathogens using multiple search platforms (MaxQuant, MSFragger, and Search GUI/PeptideShaker). The detected microbial peptides were verified using PepQuery, which analyses peptide-spectrum pairs to give statistical output for determining confident microbial peptides. Finally, a protein sequence database was generated using the list of verified microbial peptides for identification and quantitation of microbial peptides and proteins, respectively. The taxonomic analysis of the detected peptides revealed several opportunistic pathogens like *Streptococcus pneumoniae, Rhizopus microsporus, Clavispora lusitaniae*, and *Syncephalastrum racemosum* among others. Using parallel reaction monitoring (PRM), we validated a few identified microbial peptides in clinical samples. The analysis also revealed proteins belonging to species like *Pseudomonas fluorescens, Enterobacter*, and *Clostridium* to be up-regulated in severe COVID-19 samples. Thus, MS can serve as a powerful tool for untargeted detection of a wide range of microorganisms. Metaproteomic analysis in COVID-19 patients for early identification and characterisation of co-infecting microorganisms can significantly impact the diagnosis and treatment of patients.

## Introduction

The world witnessed the catastrophic pandemic of SARS-CoV-2 during the past three years, which still does not seem to be over. The pandemic wreaked havoc globally, affecting millions of lives and livelihoods, and putting immense pressure on the healthcare systems. Amidst the mayhem caused by the pandemic, secondary microbial infections added fuel to the fire. This was clearly notable from the increased reports of invasive mucormycosis post-COVID-19 cases during the second wave in India [1]. Respiratory infections disrupt the microbiota in the upper respiratory tract (URT) and put patients at risk for subsequent infections. The COVID-19 pandemic has caused a slew of microbial co-infections because of the impaired immunity and medical interventions during hospitalization. These secondary infections engendered by opportunistic pathogens have greatly hampered therapy, prognosis, and overall disease management. According to a pooled analysis of crude odds ratios for death, COVID-19 individuals who also had a co-infection had a higher mortality rate than those who did not [2]. Further, higher susceptibility to secondary infections resulted in higher mortality for severe patients as compared to non-severe patients [3]. Several cases of co-infections most commonly caused by *Staphylococcus aureus, Staphylococcus pneumoniae, Hemophilus influenzae, Acinetobacter baumanii*, and *Candida sp*. among others have been reported in COVID-19 patients across the globe [4]. Most of the pathogenic organisms found infecting COVID-19 patients around the globe were nosocomial multidrug-resistant (MDR) pathogens [5]. In the second wave of COVID-19, India witnessed a devastating endemic of mucormycosis, an aggressive and fatal fungal infection caused by the mold fungi of the order Mucorales [6]. Similarities in the clinical symptoms made the diagnosis of secondary infection challenging. This usually resulted in the development of clinical complications due to a late or incorrect diagnosis. Moreover, since most healthcare resources were focused on addressing the pandemic, medical facilities otherwise utilized for detection of secondary infections were not easily available. The prescription of broad-spectrum antibiotics also resulted in development of antimicrobial resistance [7–9]. Under such circumstances, when making diagnostic decisions for treatment are important, the detection of secondary infection-causing microorganisms is of critical importance.

Many methods are used for the clinical detection of microbial infections. This includes standard approaches like phenotype-based speciation, microscopy-based techniques, immunoassays, and advanced techniques like respiratory panels [10,11]. Although commonly employed in microbial detection, these techniques are not as efficient, especially in a pandemic situation that requires quick detection of a wide range of microbes in a large number of samples. The phenotype-based speciation approach, for example, has been found to incur many inaccuracies [12]. Similarly, there are many limitations of microscopy-based studies such as the identification of small cells, differentiation between live and dead cells, and detection amidst the noise generated by particulate matter [13]. Most importantly, the phylogenetic diversity of the microorganisms present in the study habitat is not revealed by any of these techniques [13]. Immunosorbent assays and serological tests are powerful tools for taxonomic detection but suffer from false positive detection due to the possible cross-reactivity amongst the antibodies used for detection. Several cases from Indonesia, Singapore, and Thailand reported cross-reactivity between SARS-CoV-2 and dengue antibodies, resulting in false-positive detection in dengue cases [14]. For immunosorbent assays such as ELISA, main disadvantages are that it is a labour-intensive procedure; has limited coverage of microbial detection; offers poor reproducibility, generates high rate of false positives, and is expensive [15]. The respiratory panel method is another clinical method in recent use that generates rapid results. However, this method is limited by the number of detectable microorganisms; a high cost and semi-quantitative nature of bacterial counts, and poor correlation of copy number in a large number of respiratory samples [9].

In contrast, mass spectrometry (MS)-based clinical metaproteomics can prove to be a highly useful tool, especially for detecting a wide range of microorganisms. The metaproteome is comprised of proteins that are expressed by various microorganisms found in a clinical or an environmental sample [16,17]. By analysing the expression of functional pathways and their dynamics within a microbial consortium, metaproteomics significantly improves our understanding of how the microbiome interacts with its immediate environment [17]. With the help of MS-based metaproteomic analysis of clinical isolates, one can achieve untargeted, discovery-based detection of a wide range of microorganisms such as bacteria, fungi, and viruses. In this study, MS data from nasopharyngeal swab samples of COVID-19 patients were used for metaproteomic analysis with the objective of identifying microorganisms and their potential co-infection status in disease progression.

## Methods

### Sample Information

The nasopharyngeal swab samples were obtained from Kasturba Hospital (Mumbai, India) during the first and the second wave of the pandemic in the years 2020 and 2021, respectively. These were leftover samples collected for diagnostic purposes in the hospital and were used for MS-based proteomics investigation. The nasopharyngeal swabs were collected in a viral transport medium (VTM) and the virus was heat-inactivated by incubating the samples at 65°C for 45 minutes. These samples were processed at the Proteomics Lab, Indian Institute of Technology, Bombay for MS analysis as described previously [18,19]. The MS datasets (PXD020580, PXD023016, and PXD029300) previously analysed for host proteins were analysed for their microbial composition using metaproteomic workflow (Figure 1). In addition, blank VTM without patient sample was used as control to detect microbial proteins due to environmental contamination. The VTM-containing tubes (n=4) were opened in the hospital environment and then processed, and MS data was acquired in the same manner as the actual samples.

**Figure 1:**
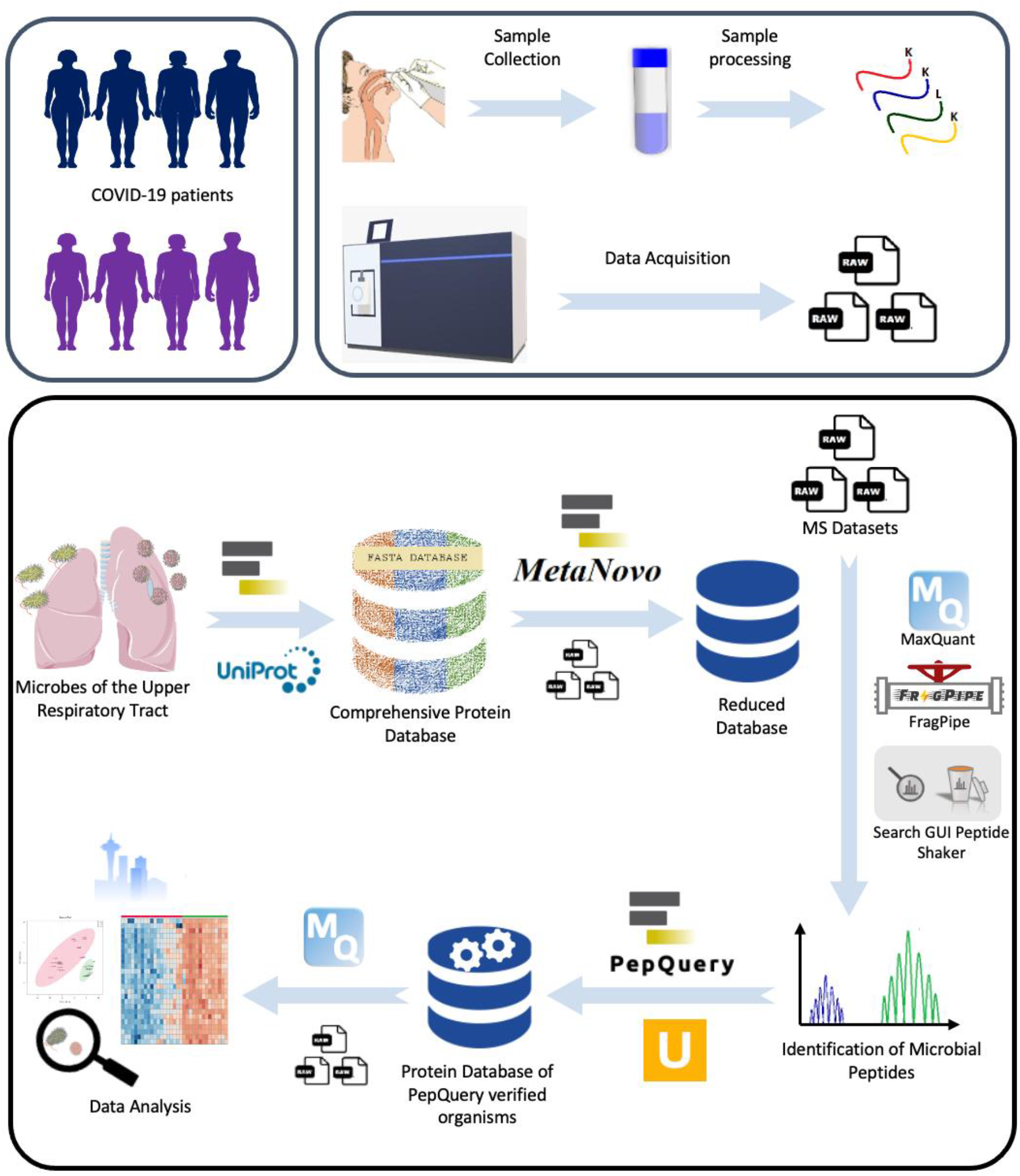
Schematic representation of the metaproteomics workflow for the analysis of the nasopharyngeal swab mass spectrometry datasets.

### Protein sequences database generation for microbial peptide detection and verification

The MS RAW files from the 2020 MS dataset (PXD020580 and PXD023016), 2021 MS dataset (PXD029300), and blank MS datasets (See section 1 “MS Datasets” in supplement) were converted into Mascot Generic Format (MGF) files using msconvert tool on Galaxy Europe server (usegalaxy.eu). A large microbial protein sequences database was generated using the Uniprot XML Downloader tool within the Galaxy platform [20] to extract sequences for known genera (33,281,528 protein sequences from 72 genera; See section 2 “Protein Sequences Database” in Supplement) and species (242,901 protein sequences from 7 species; See section 2 “Protein Sequences Database” in Supplement) from nasopharyngeal samples. These large databases were merged with the Human SwissProt database (20,361 protein sequences) and contaminant sequences database (116 protein sequences) to generate a large database of protein sequences for further processing (26,269,124 protein sequences). The MetaNovo tool [21] within Galaxy platform was used to generate a reduced protein sequences database of identified proteins. As a result, the large combined microbial database of 26,269,124 protein sequences was reduced to a compact database containing 164,023 protein sequences (for 2020 dataset search), 74,280 protein sequences (for 2021 dataset search) and 22,148 protein sequences (for Blank VTM dataset search), respectively (Supplementary Table S1, see section 2 “protein sequence databases”).

For the 2020 dataset, the 2021 dataset, and blank dataset, peptide detection was performed by using SearchGUI/Peptide Shaker [22,23] within the Galaxy platform and FragPipe [24] and MaxQuant [25] software suites. See Discovery Workflows below for details.

### Discovery Workflow 1: SearchGUI/PeptideShaker workflow

SearchGUI uses multiple search algorithms to match spectra from MS data against protein sequences search database. Search algorithms such as MS-GF+ [26], X! Tandem [27] and Comet [28] within SearchGUI were used to generate identification results from the MetaNovo-generated reduced protein sequence databases. The SearchGUI results were processed using PeptideShaker to generate outputs that were used to interpret results, in particular, to assess the confidence of the peptides detected. The identification in all discovery workflows was done as per tryptic digestion with a maximum missed cleavage of 2. Fixed modification was set to carbamidomethylation of cysteine (+57.021464 Da) and variable modification was set to oxidation of methionine (+15.994915 Da). Peptides detected were analysed for their confidence and microbial origin to generate a list of microbial peptides for the verification workflow using the PepQuery tool [29].

### Discovery Workflow 2: MSFragger workflow

MSFragger [24], a database search algorithm from the FragPipe software suite was used to search the MS files from 2020 dataset, 2021 dataset, and blank dataset against the respective Metanovo-generated reduced databases. MSFragger-detected peptides were filtered for their confidence and microbial origin to generate a list of confident peptides for the verification workflow.

### Discovery Workflow 3: MaxQuant workflow

The raw datasets were analysed using MaxQuant (version 2.0.1.0) against the respective Metanovo-generated microbial databases. The peptides output file from MaxQuant was processed wherein contaminant, human, and reverse peptides were removed, leaving just the peptides of microbial origin. The resultant microbial peptides were subjected to verification workflow using PepQuery tool.

### Microbial peptide verification workflow using PepQuery

The PepQuery tool (implemented within the Galaxy platform) was used to verify the presence of the identified confident microbial peptides from SearchGUI/PeptideShaker, MSFragger, and MaxQuant discovery workflows (see above). PepQuery (version 1) [29] was used to verify that the microbial peptides were not present in the human proteome. For PepQuery analysis, all the microbial peptides detected via discovery workflows were processed against Human UniProt, contaminants, and SARS-CoV-2 protein sequences database. PepQuery performs peptide-centric analysis of peptide-spectrum pairs involving the microbial sequences to generate statistical outputs that can be used to determine verified microbial peptides.

### Protein sequences database generation for microbial detection and quantitation

PepQuery analysis generated a master list of verified microbial peptides that was used to generate a protein sequence database for the quantitative workflow. The protein sequences of accession numbers associated with the PepQuery-verified microbial peptides were merged with Human UniProt database, contaminant sequences, and COVID-19 sequences. The resultant protein sequences database was used for the final MaxQuant quantitation workflow.

### Database search using final metaproteomics database

The MS datasets of nasopharyngeal swab samples were analysed with MaxQuant against the final reduced database. Raw files were processed using LFQ (label-free quantification) parameters with label type as “standard” and multiplicity of 1. Identification was done as per tryptic digestion with a maximum missed cleavage of 2. Fixed modification was set to carbamidomethylation of cysteine (+57.021464 Da) and variable modification was set to oxidation of methionine (+15.994915 Da). The false discovery rate (FDR) at protein and peptide level was set to 1% for reliable detection. The MaxQuant parameters used for the search are mentioned supplementary table S2 (see section 3 “MaxQuant parameters” in supplement).

The proteinGroups.txt files generated from MaxQuant analysis of the three datasets (2020, 2021, and blanks) were processed and contaminants, reverse, and human proteins were filtered out to generate a list of microbial proteins. Peptides corresponding to the microbial proteins were extracted from the peptides.txt output file of MaxQuant. The taxonomic identification of these peptides was performed using Unipept’s Metaproteomic Analysis tool [30]. Microbial peptides from all datasets (2020, 2021 and blank), that were identified at least up to genus level, were retained. These peptides were later monitored in fresh nasopharyngeal swab samples using targeted approach of parallel reaction monitoring (PRM) for validation.

### Quantitative and statistical analysis of microbial proteins

#### 2020 MS Datasets

68 nasopharyngeal swab samples from 2020 MS dataset (PXD020580 and PXD023016) were analysed using Label-Free Quantitation using MaxQuant software. Out of the 68 samples, 46 samples were classified as COVID-19 positive (labeled as “P” in our initial analysis) and 22 samples were classified as negative (labeled as “N” in our initial analysis) based on the RT-PCR results. Further, out of these 46 COVID-positive samples, 19 samples were classified as severe (labeled as “S” in subsequent analysis) and 27 were classified as non-severe (labeled as “NS” in subsequent analysis), based on clinical characteristics and oxygen supplementation.

In our initial analysis, samples were classified into two groups - COVID-19 positive (P) and COVID-19 negative samples (N). Missing values from MaxQuant outputs for LFQ intensities across the samples were checked and the protein groups with more than 50% missing values across the samples were removed. Further, data-analysis was performed using Metaboanalyst 5.0 [31]. Sample-wise K-nearest neighbour (KNN) algorithm was used for estimating missing values. The imputed missing values obtained from the previous step were then log-transformed and a correlation heatmap was plotted to determine the correlation of samples. Four samples were removed from subsequent analyses due to poor correlation.

In the subsequent analysis, the data was re-labelled as N (Negative), S (Severe) and NS (Non-Severe). In order to assess the similarity between different cohorts in our study, we used Overall Principle Component Analysis, Hierarchical Clustering, and K Means Clustering. We found significant differences between severe (S) and negative (N) patient samples. Principal component analysis (PCA) was done to specifically study the differences between severe COVID-19 patients and negative patients. Welch’s t-test was performed to identify the significant differentially expressed proteins, with a *p*-value cut-off of 0.05. Fold change analysis and T-test were performed, and the volcano plot was plotted for negative versus severe comparison. Fold change and *p*-value threshold was set to 2.0 and 0.05, respectively.

#### 2021 MS Datasets

Similar workflow was used for the analysis of swab samples from 2021 MS dataset (PXD029300). In the case of the 2021 dataset, 66 samples tested COVID-19 positive, and 9 samples tested negative using RT-PCR method. Out of the 66 positive samples, 27 were classified as severe and 39 were classified as non-severe based on clinical characteristics and oxygen supplementation. Since the proportion of negative samples was less as compared to the positive samples, only severe (S) and non-severe (N) samples were considered for comparison in the 2021 MS dataset.

### Taxonomic and Functional analysis using Unipept and BLAST-P

Unipept tool [30] was used to determine the taxonomic composition of the microbial peptide sequences. Unipept matches the peptides against the UniProt database by using the lowest common ancestor (LCA) analysis to determine the taxonomic class to which the microbial peptide can be assigned. Unipept also assigns Gene Ontology terms, EC numbers and InterPro annotation for functional analysis. The microbial peptides were also subjected to BLAST-P analysis against the NCBI non-redundant (nr) database to ascertain the microbial origin of the peptides. This verified microbial peptide panel was further used for targeted validation using parallel reaction monitoring (PRM).

### Parallel Reaction Monitoring (PRM) of identified microbial peptides in nasopharyngeal swab samples

The combined list of identified microbial peptides was used to generate an isolation list in Skyline (version 21.2.0.568) [32]. Peptides with ‘0’ missed cleavage and of 8-25 amino acids long in length were retained. The list included precursor ions (+1 and +2 charges) and b and y product ions (range “+2” to “last ion -1”) of the selected peptides.

### Sample processing for PRM

In total, 36 nasopharyngeal swab samples were processed for PRM targeted analysis. These included 18 samples from 2020 datasets (9 COVID-positive and 9 COVID-negative) and 18 samples from 2021 dataset (9 COVID-positive and 9 COVID-negative). Along with the nasopharyngeal swab samples from 2020 and 2021 datasets, four blank VTM samples were also processed for PRM as described previously [18,19]. In brief, proteins were precipitated from the samples using three different organic solvents – ethanol, isopropanol, and acetone in sample to solvent ratio of 1:3 and incubated at -20°C for 4 hrs. Protein pellets were obtained by centrifugation at 15,000g for 20 minutes at 4 °C. Supernatant was discarded and pellets were air dried followed by resuspension in urea lysis buffer (8 M Urea, 50 mM Tris pH 8.0, 75 mM NaCl, 1 mM MgCl2). Protein quantification of each precipitate was performed using Bradford assay keeping bovine serum albumin (BSA) as standard. For each sample, 15 μg of protein from each of its three precipitates was pooled, totalling to 45 μg of protein per sample. The quality of the proteins extracted was checked on SDS-PAGE for all the samples. 30 μg of protein per sample was taken ahead for in-solution digestion using trypsin followed by desalting using in-house C-18 stage tips as described [18,19]. The peptide concentration was determined using Scopes method by measuring absorbance at 205 nm and 280 nm. For PRM data acquisition, 4-5 samples per group (2020 negative, 2020 positive, 2021 negative, and 2021 positive) were pooled to a final peptide concentration of 0.25 μg/μl. All the samples were spiked in with an equal amount of heavy labelled synthetic peptide “DIFTGLIGPMK” as internal standard. The intensity of the synthetic peptide across all the samples was used to normalize the microbial peptide intensities.

### PRM Data Acquisition and Analysis

Orbitrap Fusion™ Tribrid™ Mass Spectrometer (Thermo Scientific, USA) coupled to a nanoLC system (Thermo Scientific, USA) was used for PRM experiments [33]. The isolation generated above was imported in the instrument method for PRM and used further for data acquisition. One μg of the peptides was injected using an Easy-nLC 1200 system and were separated on a 50 cm C18 analytical column (PepMap RSLC, EASY-Spray Column, Thermo Scientific, USA) at a flow rate of 300 nl/min on a linear gradient of 40 minutes. The buffer system used was 0.1% formic acid (FA) as solution A and 80% acetonitrile (ACN) in 0.1% FA as solution B. The gradient used for the chromatographic separation was the following: 2-5% buffer B for the first 5 minutes, 5-30% buffer B for the next 20 minutes, 30-48% buffer B in the next 5 minutes, 48-90% in the following 5 minutes, and then the last 5 minutes in 90% buffer B. The PRM data was analysed using Skyline (version 21.2.0.568) [32].

## Results

### Generation of reduced database and detection of microbial peptides

A major portion of the bioinformatics workflow (Figure 1) for the detection and quantitation of microbial proteins within COVID-19 patients was focused on generating a reduced database for quantitative analysis of microbial proteins. We started with a large protein sequences database (26,269,124 protein sequences) generated from known species and genera of the upper respiratory tract (see section 2 “Protein sequences database” in Supplement). MS datasets from 2020, 2021 and Blank VTM samples (see section 1 “MS Datasets” in Supplement) were used to generate a compact protein sequences database using MetaNovo software [21] within the Galaxy platform (see section 2 “Protein sequences database” in Supplement). The MetaNovo generated databases for 2020 dataset, 2021 dataset, and Blank VTM dataset were used to search against respective MS datasets using SearchGUI/PeptideShaker, MaxQuant, and FragPipe. These discovery workflows were useful to detect multiple microbial peptides. In total, 3267 microbial peptides (see section 4 “Discovery phase” in Supplement) were detected from all the MS datasets and discovery workflows. To verify these microbial peptides, we used PepQuery tool [29] resulting in detection of 2136 verified microbial peptides (see section 5 “Verification phase” in Supplement) which mapped to a total of 581 microorganisms (at least up to genera level). Out of these, 499 were identified in the patient datasets exclusively and not in blank datasets, and several of these are opportunistic pathogens. Table 1 lists some of the very common opportunistic pathogens for which at least 2 unique peptides were identified. In addition, the common opportunistic pathogens that were identified with just one unique peptide are listed in supplementary table S3 (see section 5 “Verification phase” in Supplement).

**Table 1:** List of common opportunistic pathogens detected with at least 2 unique peptides (and PepQuery verified) in nasopharyngeal swab datasets and the infections caused by them.

| Sr. No. | Microorganism | Type | Infection | Reference |
| --- | --- | --- | --- | --- |
| 1. | *Campylobacter jejuni* | Bacteria | Acute diarrhea, hemorrhagic colitis, bacteremia | [34] |
| 2. | *Streptococcus pyogenes* | Bacteria | Pharyngitis, systemic infection, toxic shock syndrome, tonsillitis, rheumatic fever, myonecrosis | [35,36] |
| 3. | *Klebsiella pneumoniae* | Bacteria | Pneumonia, bloodstream infections, meningitis | [37] |
| 4. | *Bacillus cereus* | Bacteria | Food poisoning, bacteremia | [38] |
| 5. | *Pseudomonas putida* | Bacteria | Ventilator-associated pneumonia, catheter-associated infection | [39] |
| 6. | *Bacillus subtilis* | Bacteria | Pneumonia, septicemia, endocarditis, bacteremia | [38,40] |
| 7. | *Clostridium botulinum* | Bacteria | Botulism – a flaccid paralytic disease | [41] |
| 8. | *Neisseria* | Bacteria | Pharyngitis, tonsillitis, ear infection, scarlet fever | [42] |
| 9. | *Streptococcus pneumoniae* | Bacteria | Community-acquired pneumonia, sepsis, meningitis, otitis media | [43] |
| 10. | *Serratia spp.* | Bacteria | Rare nosocomial infections | [44] |
| 11. | *Candida spp.* | Fungi | Candidiasis (superficial or invasive infection) | [45] |
| 12. | *Aspergillus spp.* | Fungi | Aspergillosis (Pulmonary infection) | [46] |
| 13. | *Rhizopus microsporus* | Fungi | Pulmonary mucormycosis | [47] |
| 14. | *Clavispora lusitaniae* | Fungi | Systemic candidiasis | [48] |
| 15. | *Syncephalastrum racemosum* | Fungi | Mucormycosis | [49] |
| 16. | *Cryptococcus neoformans* | Fungi | Pulmonary infection, meningoencephalitis | [50] |
| 17. | *Fusarium spp.* | Fungi | Superficial, locally invasive, or disseminated infections (keratitis, onychomycosis, erythematous) | [51,52] |

The verified microbial peptides were used to generate a list of accession numbers to construct a verified microbial proteins database. Verified microbial protein sequences (1904 protein sequences) were merged with Human UniProt database and contaminants to generate a verified microbial database (103,389 protein sequences, see section 6 “Final Search Database Generation” in Supplement) for further quantitative analysis of microbial proteins using MaxQuant platform (see section 7 “Quantitation Outputs From MaxQuant Search” in Supplement).

### Database search using final metaproteomics database

To investigate alterations in the microbial proteins, the MS datasets were analysed using MaxQuant against the final reduced database. The MaxQuant searches using the verified microbial database detected 161, 3213, and 1984 proteins in Blank, 2020, and 2021 MS datasets, respectively (Table 2). Out of these, 93 proteins (Blank MS dataset), 387 proteins (2020 MS dataset) and 253 proteins (2021 MS dataset) were of microbial origin (Table 2). Microbial peptides were subjected to Unipept analysis for all datasets and assigned to genera using the lowest common ancestor method. Figure 2 shows the overlap amongst microbial proteins, microbial peptides, and genera in 2020, 2021, and blank MS datasets. The microbial peptides associated with at least up to genera level were combined and monitored in fresh nasopharyngeal swab samples by PRM.

**Table 2:** Table showing the total number of proteins, corresponding peptides, spectra and mapped microbes (at least up to genus level) after removing human, reverse, and contaminant proteins. The numbers in the parenthesis represent the same corresponding to the mapped microbes.

| Dataset | Proteins | Peptides | Spectra | Peptides assigned at the genus level |
| --- | --- | --- | --- | --- |
| Blank | 161 (93) | 269 (96) | 625 (332) | 56 |
| 2020 | 4213 (387) | 28258 (527) | 102797 (8821) | 259 |
| 2021 | 1984 (253) | 10808 (347) | 63505 (7131) | 143 |

**Figure 2:**
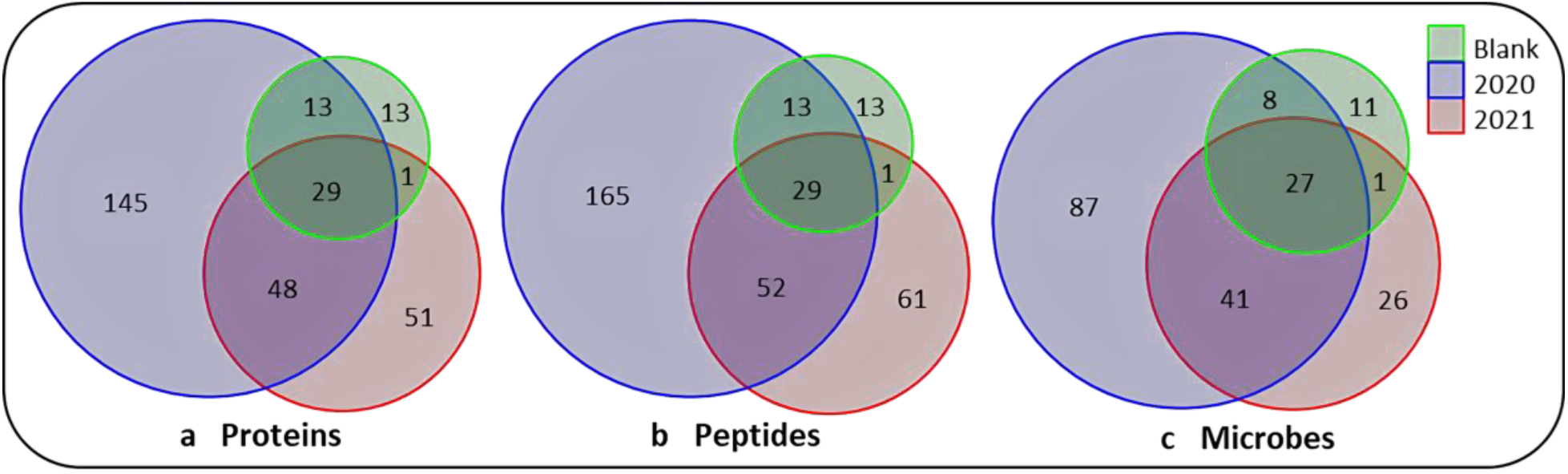
Venn diagrams representing the number of (a) filtered proteins (only microbial), (b) corresponding peptides and (c) mapped microbes (at least up to genera level).

### Quantitative and statistical analysis of microbial proteins

The MaxQuant microbial protein outputs from MS datasets were subjected to quantitative analysis. For the 2020 dataset (PXD020580 and PXD023016), the initial analysis between positive (P) and negative (N) groups using hierarchical clustering heatmap and PCA indicated that the data had additional subgroups in the data (Supplementary Figure S1, see section 7 “Quantitation outputs from MaxQuant search” in supplement). As a result, positive samples were sub-grouped as severe (S) and non-severe (NS). These new groups were subjected to hierarchical clustering and PCA, which indicated that the COVID-19 severe and negative samples were segregating, and the non-severe samples were ubiquitously spread (Supplementary Figure S2, see section 7 “Quantitation outputs from MaxQuant search” in supplement). Thus we decided to take forward the negative samples (n=22) and the severe samples (n=19) for analysis. One striking observation was the erratic behaviour of 5 severe samples, hinting towards the possibility of them being potential outliers (Supplementary Figure S3a, see section 7 “Quantitation outputs from MaxQuant search” in supplement). Subsequent Random Forest algorithm analysis predicted that five severe samples (S23, S29, S165, S155, and S167) were potential outliers (Supplementary Figure S3b, see section 7 “Quantitation outputs from MaxQuant search” in supplement) and, thus, were excluded from further analyses.

The microbial proteins from the 22 negative samples and 14 severe samples demonstrated a very clear separation in PCA plot (Figure 3a). Further analysis revealed 60 up-regulated and 5 down-regulated microbial proteins with a fold change value greater than 2 and p-value less than 0.05 (Figure 3b and Supplementary Table S4, see section 7 “Quantitation outputs from MaxQuant search” in supplement). The top 25 significant proteins that segregated negative and severe are depicted using hierarchical clustering-based heatmap in figure 3c. The top six statistically-significant differentially up-regulated microbial proteins in severe group included proteins from potential pathogens such as *Pseudomonas fluorescens* (A0A5E7DY38), *Clostridium* sp. (R7B9V3), *Penicillium steckii* (A0A1V6T7K4), *Enterobacter* (A0A0X4EHT2), *Fusarium vanettenii* (C7ZQE2), and *Siminovitchia fortis* (A0A443IKI4) (Figure 3b).

**Figure 3:**
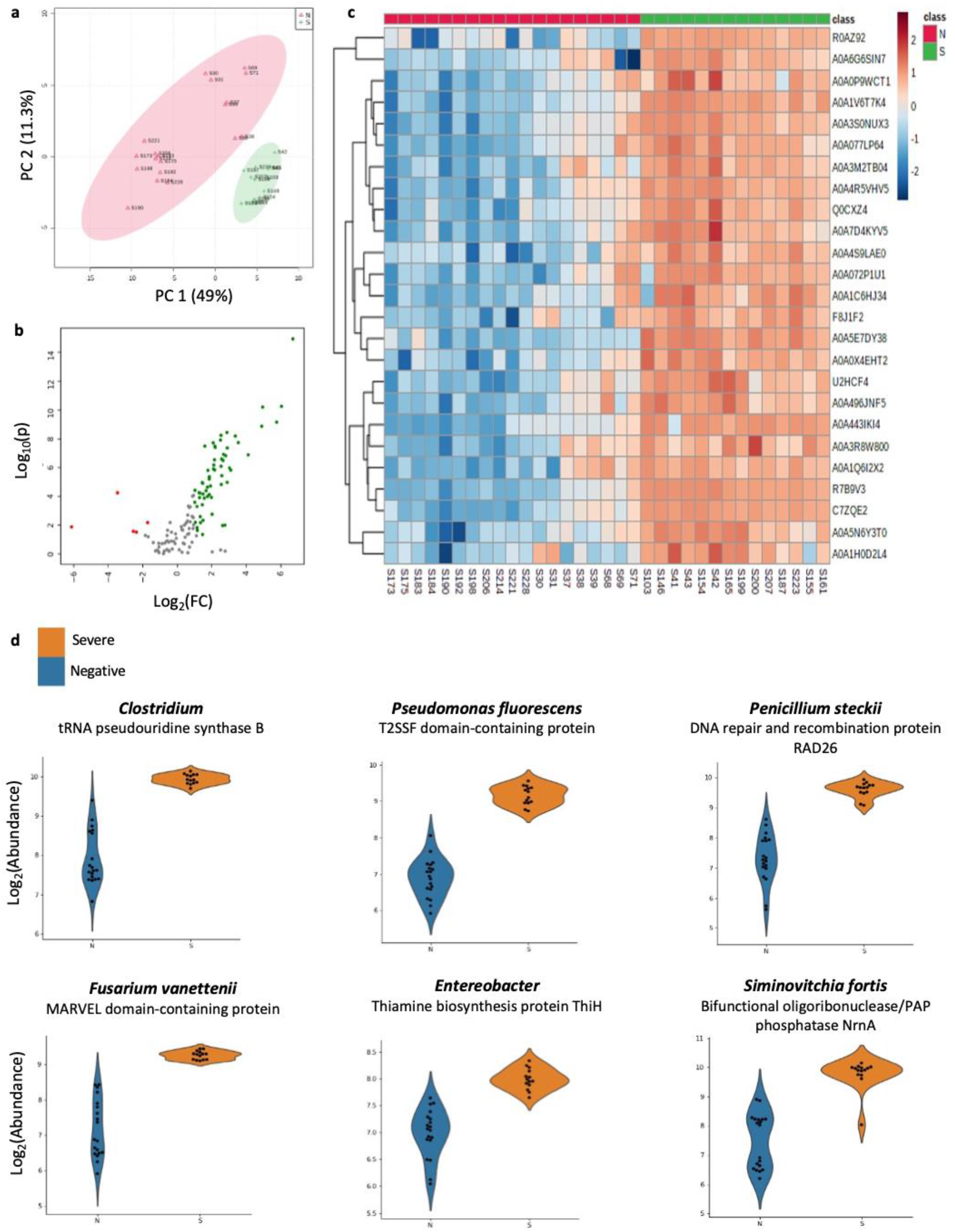
Metaproteomic analysis of COVID-19 severe and COVID-19 negative patients from first wave of COVID-19 in India (2020): (a) PCA plot showing segregation of negative and severe COVID-19 samples. (b) Volcano plot showing differentially expressed microbial proteins with fold change>2 and p-value cut-off of 0.05. (c) Heatmap of top 25 differentially expressed microbial proteins in severe vs negative samples. (d) Violin plots of the top six differentially expressed proteins in severe vs. negative samples.

Similar statistical analysis was also performed for the 2021 MS dataset. However, we did not observe any statistically significant differences. Hierarchical clustering and PCA exhibited a clear overlap between severe and non-severe cohorts from 2021 MS datasets (Supplementary figures S4a and S4b, see section 7 “Quantitation outputs from MaxQuant search” in supplement). Due to lack of any meaningful biological signals, we did not pursue further analysis on the 2021 MS dataset.

### Validation of microbial peptides by parallel reaction monitoring (PRM)

The microbial peptides that were detected in the MaxQuant search and assigned to the genus level were monitored in nasopharyngeal swab samples using PRM. The RAW files generated from the PRM analysis were imported into Skyline software for refining and peak-filtering. PROSIT and LFQ generated libraries were used for spectral matching to gain confidence in peptide detection. Thirteen microbial peptides were detected that showed good intensity peaks in patient samples with no or low intensity peak in blank VTMs. These peptides belong to microorganisms such as *Bacillus sp*., *Sphingobacterium sp*., *Staphylococcus sp*., *Aspergillus sp*., *Penicillium steckii, Clostridium sp*., *Lactobacillus sp*., *Burkholderia cenocepacia, Fusarium sambucinum*, and *Actinomyces bowdenii* (Figure 4 and supplementary figure S5, section 8 “parallel reaction monitoring of microbial peptides” in supplement). All the reported peptides have a library-peptide match score of greater than 0.7.

**Figure 4:**
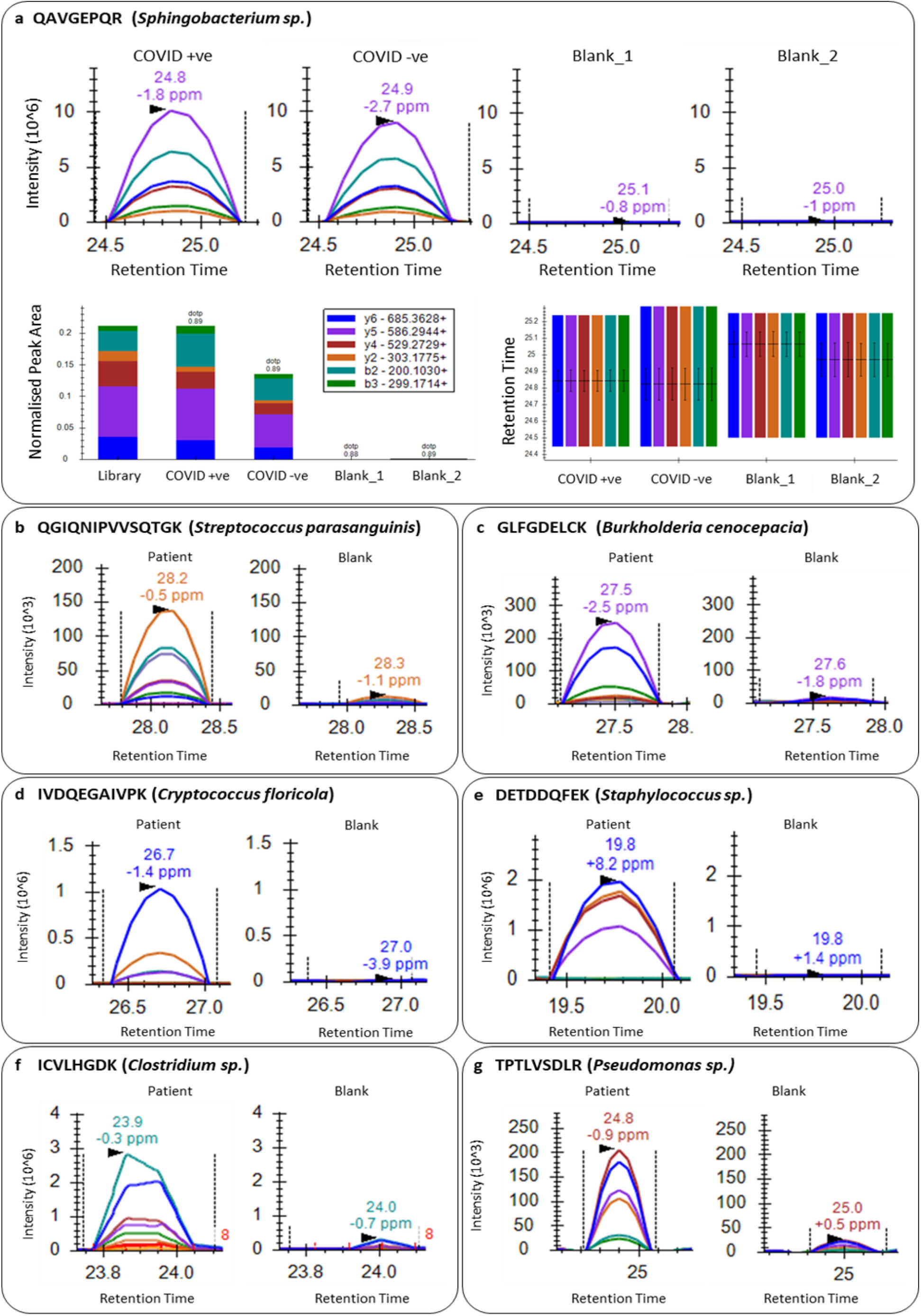
Detection of microbial peptides in nasopharyngeal swab samples by PRM: (a) QAVGEPQR (*Sphingobacterium sp*.) in COVID-positive, COVID-negative, and blank samples (top panel); normalized peak area (bottom left) and retention time (bottom right) in individual samples. (b), (c), (d), (e), (f), and (g) show peak intensities for peptides QGIQNIPVVSQTGK, GLFGDELCK, IVDQEGAIVPK, DETDDQFEK, ICVLHGDK, and TPTLVSDLR, respectively in patient sample vs blank.

## Discussion

In this clinical metaproteomics investigation, we have utilized MS-based proteomics data from nasopharyngeal swabs of COVID-19 patients to investigate the microbial composition in the clinical samples. We have optimized a bioinformatics workflow [53] to generate a concise and specific protein database from a large and comprehensive protein database to facilitate fast and efficient database search to study the metaproteome of the nasopharyngeal swab samples. We compiled protein FASTA sequences of relevant microbes of the URT to generate a comprehensive metaproteomic database. This huge database, then, was reduced to a smaller one using Metanovo, which uses *de novo* sequence tag matching along with an algorithm for optimization of the UniProt database to generate customized protein sequences database [21].

We report the detection of microbial peptides and proteins in nasopharyngeal swab samples using this reduced metaproteomic database. To maximize the identification of microbial peptides, we used three different platforms (SearchGUI/Peptide Shaker, MaxQuant, and MS Fragger). Microbial peptides detected using all three platforms were further verified using PepQuery. With this approach, we identified peptides from several opportunistic bacterial and fungal pathogens like *Bacillus subtilis, Clostridiuim botulinum, Streptococcus pneumoniae, Klebsiella pneumoniae, Candida albicans*, and *Syncephalastrum racemosum* among many others (Table 1). *S. pneumoniae, H. influenzae*, and *K. pneumoniae* are among the most commonly reported secondary infections reported in COVID-19 cases globally [4,9,53–55]. We also detected multiple peptides from fungal species of the order Mucorales, e.g., *Rhizopus microspores* and *Syncephalastrum racemosum*. These are known to cause mucormycosis, a rare but potentially fatal disease reported in severe and recovering COVID-19 patients during the second wave of the pandemic in India [6,56].

We also validated a few of the identified microbial peptides in fresh nasopharyngeal swab samples using PRM. Of these, peptides belonging to *Burkholderia cenocepacia, Cryptococcus floricola, Streptococcus parasanguinis, Clostridium sp*., *Staphylococcus sp*., *and Sphingobacterium sp*., are worth noting as they are known causative agents of various human infections. Notably, *B. cenocepacia* is a biofilm-producing opportunistic human pathogen responsible for causing fatal nosocomial lung infections in immunocompromised patients, especially those with cystic fibrosis [57]. Moreover, *S. parasanguinis* is a commensal bacterium inhabiting multiple locations including oral cavity and intestines in humans, and is involved in causing dental plaques and occasionally infective endocarditis [58,59]. Interestingly, we also found a peptide belonging to *Glaesserella parasuis*, a very common respiratory disease-causing bacterium of pigs [60]. It is a commensal bacterium in the porcine nasal cavity and can cause serious systemic infection, characterized by arthritis, meningitis, and polyserositis [61]. Very recently, *G. parasuis* DNA has been identified for the first time in periodontal tissues in adult humans and has been implicated in triggering rheumatoid arthritis through molecular mimicry [62]. Our results too add evidence to the presence of *G. parasuis* in humans, with the potential of it being an emerging human pathogen. Surprisingly, peptides belonging to *Actinomyces bowdenii* (causes infection in cats and dogs [63]), *Penicillium steckii* (a marine endozoic fungus [64]), and *Fusarium sps*. (a mycotoxigenic plant pathogen [65]) were also detected in patient samples. These have never been reported to inhabit or infect humans and further studies are required to substantiate these findings.

Further, we report dysregulation of some microbial proteins in severe COVID-19 patients as compared to negative patients from the 2020 MS dataset. The top six proteins that were significantly up-regulated in the severe group belong to microbes like *Pseudomonas fluorescens, Clostridium* sp., *Penicillium steckii, Enterobacter, Fusarium vanettenii*, and *Siminovitchia fortis*. Surprisingly 3 of them, i.e., *Penicillium steckii, Fusarium vanettenii*, and *Siminovitchia fortis*, have never been reported to infect humans. *Pseudomonas fluorescens*, which is a Gram-negative commensal bacterium of human digestive tract, is mostly non-pathogenic, but has been reported to be identified in respiratory samples and to cause opportunistic bacteremia [66]. Additionally, several species of the genera *Enterobacter* and *Clostridium* are very common opportunistic pathogens that can cause life-threatening infections in immunocompromised individuals. Many proteins belonging to other opportunistic pathogens like *Bacteroides fragilis, Proteus vulgaris, Helicobacter pylori*, and *Bacillus cereus* were also upregulated in severe group (Supplementary table S4, see section 7 “Quantitation outputs from MaxQuant search” in supplement) signifying the higher possibility of severe patients developing secondary infections. We did not observe any statistically significant differential expression of microbial proteins in 2021 datasets. This could be due to the comparison between non-severe and severe groups as opposed to the severe vs negative comparison of the 2020 dataset as the number of negative samples was limited in 2021 dataset. This also corroborates with the observation from the 2020 dataset principal component analysis (Supplementary figure S2, see section 7 “Quantitation outputs from MaxQuant search” in supplement) wherein the non-severe group was ubiquitously spread and did not cluster well.

Apart from the differential expression of some microbial proteins observed in the severe group as compared to the negative group, there was no significant difference in the levels of several other microbial peptides and proteins. Peptides belonging to many common opportunistic pathogens were detected in individuals from both positive and negative datasets. This is not surprising as these microorganisms are a part of the natural microbial flora in humans but are more likely to manifest as infections in immunocompromised patients. The presence of a potential pathogen serves as a pathogenicity indicator and would require further clinical tests to verify this. A limitation of the study is that all the samples were collected from hospitalized patients. Even the negative samples were from patients who had tested negative after recovering from COVID-19 thus being exposed to the same hospital environment as the positive patients. Another limitation of our study is that the nasopharyngeal swab samples were originally collected for diagnostic purposes. The left-over samples, after diagnostic testing, were utilized and processed to study the host response using mass-spectrometry-based proteomics [18] and the same MS data was used for this metaproteomics investigation.

Despite these limitations, we have established a robust workflow to study the metaproteome of clinical samples from MS data. We have identified some opportunistic pathogens from diagnostic samples of COVID-19 patients, which can be monitored to stratify high-risk patients. We believe that clinical metaproteomics can serve as an excellent tool to aid untargeted diagnosis and, thereby, better management of secondary infections, especially in the face of pandemic such as that of COVID-19. Identification of novel and emerging pathogens and antibiotic-resistant microorganisms from clinical specimens can complement clinical testing and assist in making informed judgements. Furthermore, a similar workflow can be utilised for waste-water microbiome surveillance to track community-wide emergence and transmission of infectious agents and their variants [67]. This can be crucial in case of outbreaks and pandemics, where epidemiological trends can be identified earlier than clinical case reporting, thereby helping in preparedness for the disease.

## Data Availability

All the information regarding the data presented in this article is mentioned in the supporting information. The raw files for the 2020 MS dataset can be accessed via PRIDE ID: PXD020580 and PXD023016. The raw files for the 2021 MS dataset can be accessed via PRIDE ID: PXD029300. The raw files for the blank dataset have been uploaded on Galaxy Europe server (usegalaxy.eu) and can be accessed via link: https://usegalaxy.eu/u/galaxyp/h/inputiitbombayblanks. The protein sequence databases used for this study have also been uploaded on Galaxy Europe server and can be accessed via link: https://usegalaxy.eu/u/galaxyp/h/covid-metaproteomics-databases-2020-2021. The targeted proteomics data has been deposited in Proteome Xchange Consortium and can be accessed via the link: https://panoramaweb.org/COVID19metaproteomics.url.

## Acknowledgements

Our deepest gratitude to Kasturba Hospital for Infectious Diseases for allowing us to collect COVID-19 samples and for providing the clinical information of the patients. The study was supported through Science and Engineering Research Board (SERB), Department of Science & Technology, Ministry of Science and Technology, Government of India (SB/S1/Covid-2/2020), and a special COVID-19 seed grant (RD/0520-IRCCHC0-006) from IRCC, IIT Bombay to SS. The authors want to thank MERCK-COE (DO/2021-MLSP) for their extended support. MASSFIITB (Mass Spectrometry Facility, IIT Bombay) from the Department of Biotechnology (BT/PR13114/INF/22/206/2015) is gratefully acknowledged for MS-based proteomics work. The authors would also like to acknowledge the Thermo Fisher Scientific engineers and application scientists for their constant support to the MASSFIIT facility, IIT Bombay during the pandemic. S.B. acknowledges the CSIR funding for PhD. D.B. is supported by MHRD for M.Tech.

## Author Contributions

P.J., S.S. and S.B. conceived and designed the project. S.M., P.J., A.R., and J.J. performed the data analysis involving Metanovo, Galaxy workflows, and Fragpipe. A.G., S.B., and D.B. performed the data analysis involving MaxQuant and further taxonomic analysis. S.B. and D.B. conducted the PRM experiments. S.B., S.M., D.B., A.G., and P.J. have drafted the original manuscript. P.J., S.S, S.B., S.M., and T.G. have edited and reviewed the manuscript. All the authors have approved the final version of the manuscript.

