## Supplementary Data for "Metaproteomic analysis of nasopharyngeal swab samples to identify microbial peptides and potential co-infection status in COVID-19 patients"

**Section 1. MS DATASETS:**

1) BLANKS: https://usegalaxy.eu/u/galaxyp/h/inputiitbombayblanks
2) 2020 Mass Spectrometry Datasets:
   http://proteomecentral.proteomexchange.org/cgi/GetDataset?ID=PXD020580
   http://proteomecentral.proteomexchange.org/cgi/GetDataset?ID=PXD023016
3) 2021 Mass Spectrometry Dataset:
   Project Name: Proteomic analysis of swab samples from COVID-19 patients in the second wave
   Project accession: PXD029300

**Section 2. PROTEIN SEQUENCE DATABASES:**

History: https://usegalaxy.eu/u/galaxyp/h/covid-metaproteomics-databases-2020-2021

**Table S1**. Number of proteins sequences in each database.

| | |
|---|---|
| **Large database** | 26,26,9124 sequences |
| **Discovery-based database** | 164,023 sequences (2020) |
| | 74,280 sequences (2021) |
| | 22,148 sequences (Blank) |
| **Final reduced database** | 103,389 sequences |

**Section 3. MAXQUANT PARAMETERS**

**Table S2**. MaxQuant parameters for database search using metaproteomics database

| **Parameter** | **Value** |
|---|---|
| Version | 2.0.1.0 |
| User name | Admin |
| Machine name | ADMIN-PC |
| Include contaminants | TRUE |
| PSM FDR | 0.01 |

| | |
|---|---|
| PSM FDR Crosslink | 0.01 |
| Peptide FDR | 0.01 |
| Protein FDR | 0.01 |
| Site FDR | 0.01 |
| Min. peptide Length | 8 |
| Min. score for unmodified peptides | 0 |
| Min. score for modified peptides | 40 |
| Min. delta score for unmodified peptides | 0 |
| Min. delta score for modified peptides | 6 |
| Min. unique peptides | 1 |
| Min. razor peptides | 1 |
| Min. peptides | 1 |
| Use only unmodified peptides | TRUE |
| Modifications included in protein quantification | Carbamidomethyl (C), Oxidation (M),Acetyl (Protein N-term) |
| Peptides used for protein quantification | Razor |
| Discard unmodified counterpart peptides | TRUE |
| Label min. ratio count | 2 |
| Use delta score | FALSE |
| iBAQ | FALSE |
| Instrument Type | Orbitrap |
| Match between runs | TRUE |
| Matching time window [min] | 0.7 |
| Match ion mobility window [indices] | 0.05 |
| Alignment time window [min] | 10 |
| Alignment ion mobility window [indices] | 1 |
| Find dependent peptides | FALSE |
| Decoy mode | revert |
| Include contaminants | TRUE |
| Advanced ratios | TRUE |
| Second peptides | TRUE |

| | |
|---|---|
| Stabilize large LFQ ratios | TRUE |
| Separate LFQ in parameter groups | FALSE |
| Require MS/MS for LFQ comparisons | TRUE |
| Main search max. combinations | 200 |
| Advanced site intensities | TRUE |
| Max. peptide mass [Da] | 4600 |
| Min. peptide length for unspecific search | 8 |
| Max. peptide length for unspecific search | 25 |
| Razor protein FDR | TRUE |
| Disable MD5 | FALSE |
| Max mods in site table | 3 |
| Match unidentified features | FALSE |
| Evaluate variant peptides separately | TRUE |
| Variation mode | None |
| MS/MS tol. (FTMS) | 20 ppm |
| Top MS/MS peaks per Da interval. (FTMS) | 12 |
| Da Interval. (FTMS) | 100 |
| MS/MS deisotoping (FTMS) | TRUE |
| MS/MS deisotoping tolerance (FTMS) | 7 ppm |
| MS/MS tol. (ITMS) | 0.5 Da |
| Top MS/MS peaks per Da interval. (ITMS) | 8 |
| Da interval. (ITMS) | 100 |
| MS/MS deisotoping (ITMS) | FALSE |
| MS/MS deisotoping tolerance (ITMS) | 0.15 Da |
| MS/MS ammonia loss (ITMS) | TRUE |
| MS/MS dependent losses (ITMS) | TRUE |
| MS/MS recalibration (ITMS) | FALSE |
| MS/MS tol. (Unknown) | 0.5 Da |
| Top MS/MS peaks per Da interval. (Unknown) | 8 |

| | |
|---|---|
| Da interval. (Unknown) | 100 |

**Section 4. DISCOVERY PHASE:**

1) SearchGUI/PeptideShaker Discovery Workflow:
   History: https://usegalaxy.eu/u/galaxyp/h/covidsgpsmp-searches
   Workflow: https://usegalaxy.eu/u/galaxyp/w/searchguipeptideshaker-mp-covid

2) FragPipe Discovery Workflow:
   History: https://usegalaxy.eu/u/galaxyp/h/covid-fragpipe-metaproteomics-searches

3) MaxQuant Discovery Workflow:
   History: https://usegalaxy.eu/u/galaxyp/h/covid-maxquant-mp-searches

4) Microbial peptides from Discovery Workflow:
   History: https://usegalaxy.eu/u/galaxyp/h/microbial-peptides-from-discovery-searches-1

**Section 5. VERIFICATION PHASE:**

History: https://usegalaxy.eu/u/galaxyp/h/covid-pepquery-mp-blank-2020-2021

**Table S3**: List of common opportunistic pathogens detected with 1 unique, and PepQuery verified peptide in nasopharyngeal swab datasets and the infections caused by them. This table is an extension of the Table 1 in the main manuscript that includes list of organisms with at least two unique, and PepQuery verified peptides, each.

| Sr. No. | Microorganism | Type | Infection | Reference |
|---|---|---|---|---|
| 1 | *Haemophilus influenzae* | Bacteria | Respiratory tract infection, pneumonia | 1 |
| 2 | Prevotella intermedia | Bacteria | Periodontal infection | 2 |
| 3 | *Bacteroides pyogenes* | Bacteria | Bacteremia | 3 |
| 4 | *Ralstonia insidiosa* | Bacteria | Nosocomial infections | 4 |

| | | | | |
|---|---|---|---|---|
| 5 | *Ralstonia pickettii* | Bacteria | Nosocomial infections | 4 |
| 6 | *Acinetobacter johnsonii* | Bacteria | Nosocomial infections | 5 |
| 7 | *Lichtheimia ramosa* | Fungi | Pulmonary, rhino-cerebral, CNS, or cutaneous infections (mucormycosis) | 6 |


**Section 6. FINAL SEARCH DATABASE GENERATION:**

History:
https://usegalaxy.eu/u/galaxyp/h/finalsearchdatabasegeneration

Workflows:
Accession retrieval from MaxQuant Searches:
https://usegalaxy.eu/u/galaxyp/w/maxquant-accession-numbers-of-pepquery-verified-peptides
Accession retrieval from SearchGUI/PeptideShaker Searches:
https://usegalaxy.eu/u/galaxyp/w/imported-sgps-accession-numbers-of-pepquery-verified-peptides
Accession retrieval from FragPipe Searches:
https://usegalaxy.eu/u/galaxyp/w/fragpipe-accession-numbers-of-pepquery-verified-peptides

**Section 7. QUANTITATION OUTPUTS FROM MAXQUANT SEARCH:**

MaxQuant and Unipept outputs for blank, 2020 and 2021
History: https://usegalaxy.eu/u/galaxyp/h/maxquantunipeptblank-2020-2021

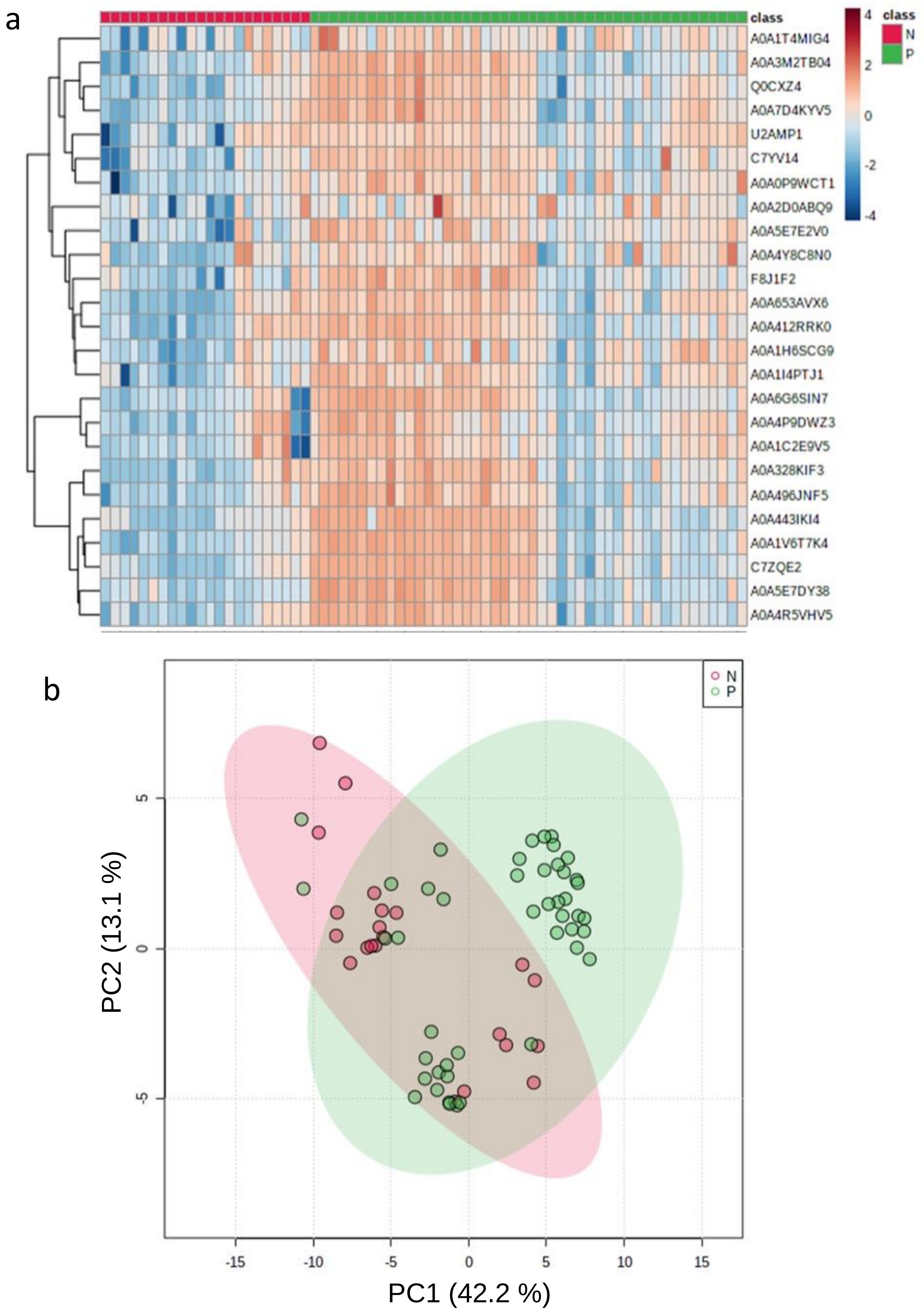


**Figure S1.** Metaproteomic analysis of COVID-19 positive (P) and COVID-19 negative (N) patients from first wave of COVID-19 in India (2020): (a) Hierarchical clustering heatmap and (b) PCA plot for positive and negative group comparison.

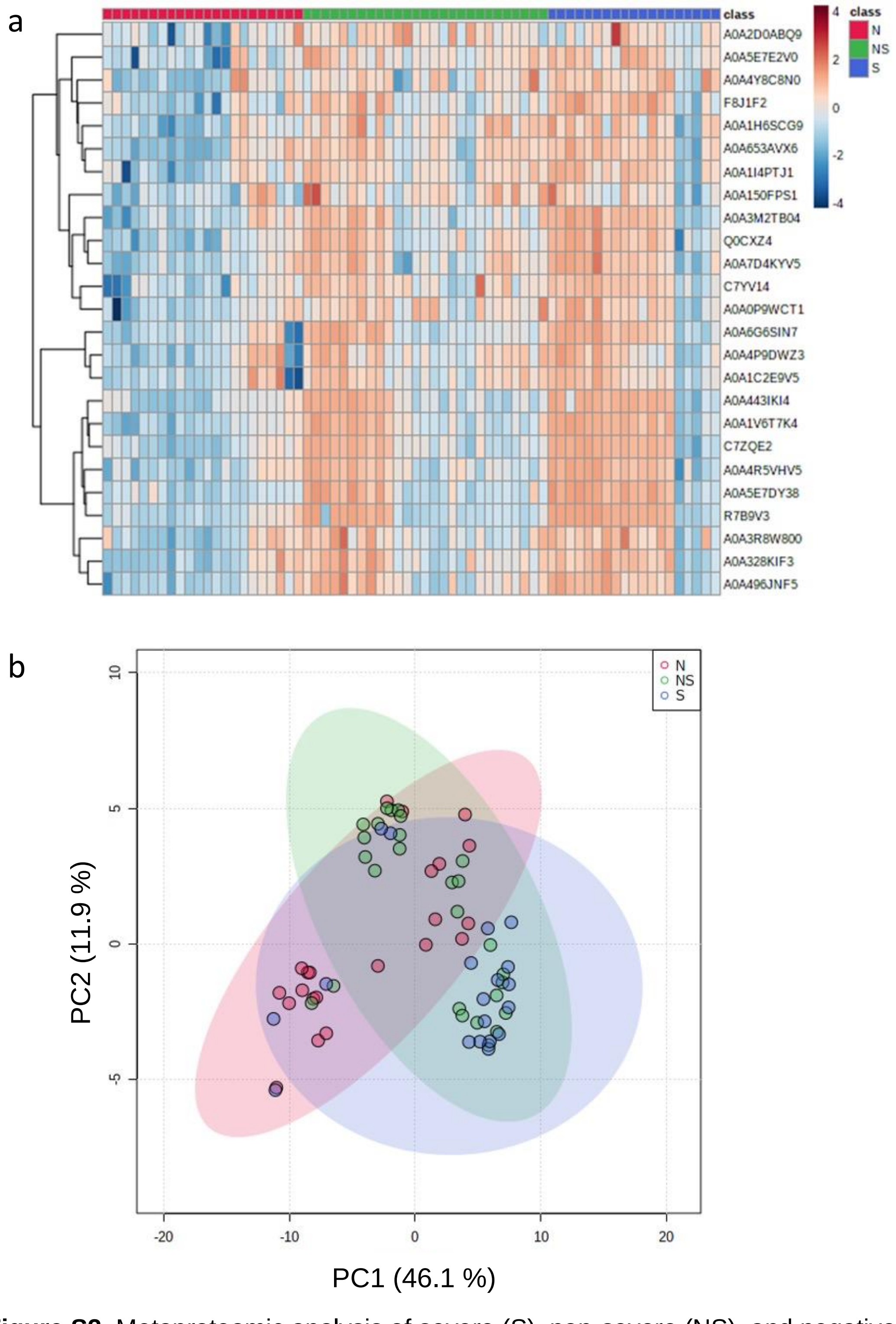


**Figure S2.** Metaproteomic analysis of severe (S), non-severe (NS), and negative (N) COVID-19 patients from first wave of COVID-19 in India (2020): (a) Hierarchical clustering heatmap and (b) PCA plot for negative, non-severe, and severe group comparison.

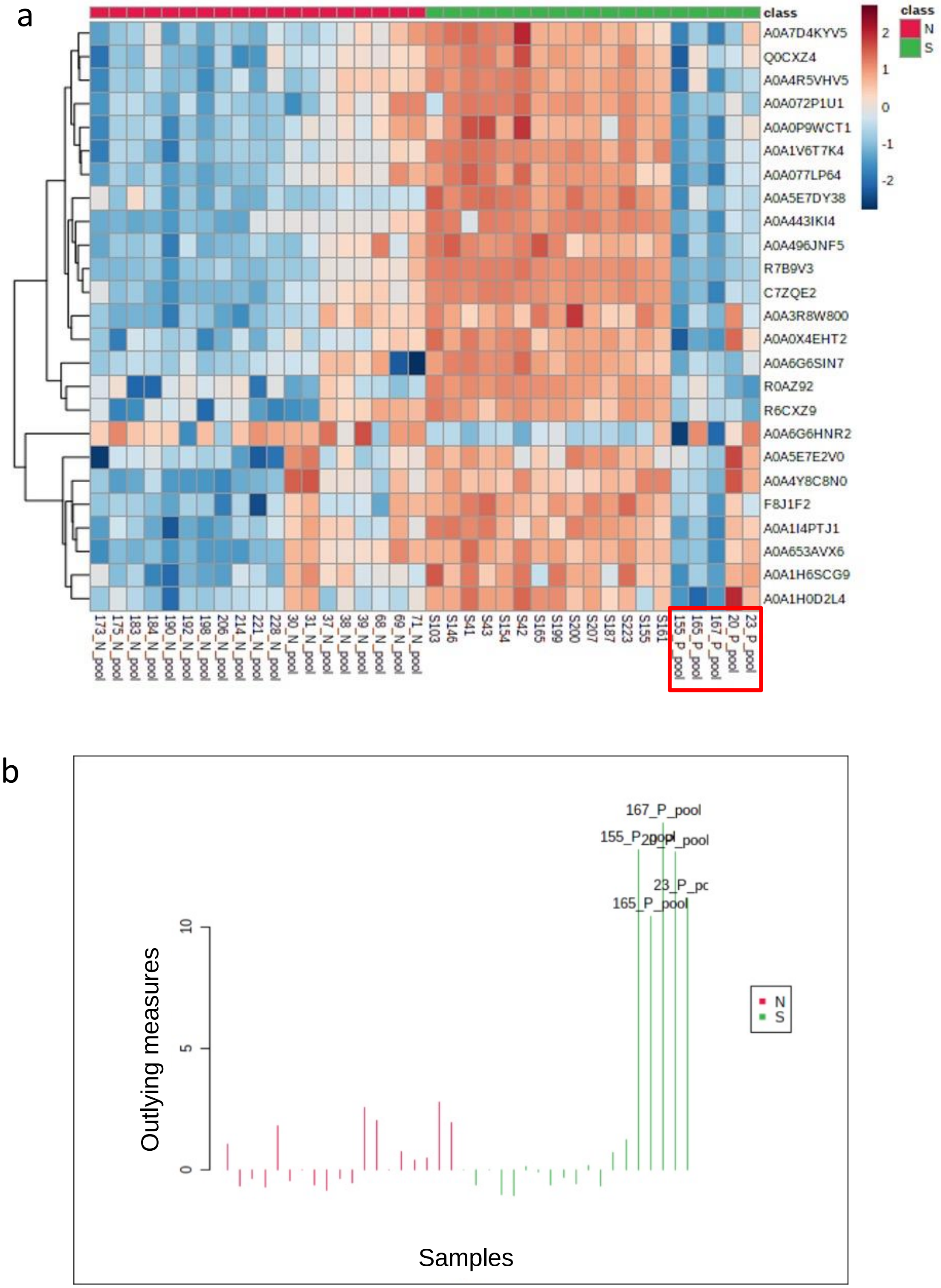


**Figure S3:** Identification of outliers in 2020 dataset: **(a)** Five positive samples showing erratic behaviour in hierarchical clustering **(b)** Outlier detection method based on random forest algorithm predicts the same five samples as potential outliers.

**Table S4**: List of top 65 differentially expressed proteins in severe group as compared to the negative group

| Uniprot ID | Protein names | Organism | Fold Change (FC) | log2(FC) | Raw p-value | -log10(pval) |
|---|---|---|---|---|---|---|
| A0A5E7DY38 | T2SSF domain-containing protein | *Pseudomonas fluorescens* | 104.94 | 6.71 | 1.17E-15 | 14.93 |
| A0A1V6T7K4 | DNA repair and recombination protein RAD26 | *Penicillium steckii* | 66.05 | 6.05 | 5.65E-11 | 10.25 |
| R7B9V3 | tRNA pseudouridine synthase B (EC 5.4.99.25) | *Clostridium* sp. CAG:505 | 31.06 | 4.96 | 6.35E-11 | 10.20 |
| A0A443IKI4 | Bifunctional oligoribonuclease/PAP phosphatase NrnA | *Siminovitchia fortis* | 54.33 | 5.76 | 6.98E-10 | 9.16 |
| C7ZQE2 | MARVEL domain-containing protein | *Fusarium vanettenii* | 29.98 | 4.91 | 1.35E-09 | 8.87 |
| A0A0X4EHT2 | Thiamine biosynthesis protein ThiH | *Enterobacter genomosp. O* | 7.37 | 2.88 | 3.63E-09 | 8.44 |
| U2HCF4 | IPT/TIG domain-containing protein | *Sphingobacterium paucimobilis* HER1398 | 5.80 | 2.54 | 5.72E-09 | 8.24 |
| Q0CXZ4 | S-adenosyl-L-methionine-dependent methyltransferase | *Aspergillus terreus* (strain NIH 2624 / FGSC A1156) | 10.41 | 3.38 | 6.41E-09 | 8.19 |
| A0A7D4KYV5 | DUF4178 domain-containing protein | *Bacteroides fragilis* | 5.70 | 2.51 | 1.21E-08 | 7.92 |
| A0A4R5VHV5 | Haloacid dehalogenase-like hydrolase | *Bacillus salipaludis* | 11.72 | 3.55 | 1.82E-08 | 7.74 |
| A0A496JNF5 | - | - | 4.25 | 2.09 | 1.88E-08 | 7.72 |
| A0A077LP64 | Histidine kinase (EC 2.7.13.3) | *Pseudomonas sp*. StFLB209 | 3.05 | 1.61 | 3.18E-08 | 7.50 |
| A0A3S0NUX3 | - | - | 4.55 | 2.18 | 3.21E-08 | 7.49 |

| | | | | | | |
|---|---|---|---|---|---|---|
| A0A3M2TB04 | Oxidoreductase | *Aspergillus sp*. HF37 | 4.62 | 2.21 | 3.86E-08 | 7.41 |
| A0A5N6Y3T0 | PHD-type domain-containing protein | *Aspergillus arachidicola* | 7.46 | 2.90 | 4.29E-08 | 7.37 |
| F8J1F2 | Phage protein | *Lactobacillus* phage JCL1032 | 6.43 | 2.69 | 1.25E-07 | 6.90 |
| R0AZ92 | Stage 0 sporulation protein A homolog | *Enterocloster bolteae* 90B8 | 17.44 | 4.12 | 1.35E-07 | 6.87 |
| A0A4S9LAE0 | MBOAT_2 domain-containing protein | *Aureobasidium pullulans* (Black yeast) (*Pullularia pullulans*) | 6.99 | 2.81 | 1.57E-07 | 6.81 |
| A0A072P1U1 | PilX_N domain-containing protein | *Schinkia azotoformans* MEV2011 | 5.99 | 2.58 | 2.63E-07 | 6.58 |
| A0A3R8W800 | DUF4148 domain-containing protein | *Pseudomonas sp*. p106 | 4.48 | 2.16 | 3.01E-07 | 6.52 |
| A0A0P9WCT1 | DUF1616 domain-containing protein | *Pseudomonas coronafaciens pv. oryzae* | 6.32 | 2.66 | 3.32E-07 | 6.48 |
| A0A1H0D2L4 | Spore germination protein KC | *Bacillus sp*. OK048 | 4.37 | 2.13 | 6.89E-07 | 6.16 |
| A0A6G6SIN7 | Basal-body rod modification protein FlgD | *Proteus vulgaris* | 5.83 | 2.54 | 8.33E-07 | 6.08 |
| A0A1C6HJ34 | Uncharacterized protein conserved in bacteria | uncultured *Clostridium* sp | 8.21 | 3.04 | 1.09E-06 | 5.96 |
| A0A1Q6I2X2 | Glyco_trans_2-like domain-containing protein | *Bacteroides uniformis* | 4.32 | 2.11 | 1.30E-06 | 5.89 |
| A0A1I4PTJ1 | Arsenate reductase | Pseudomonas yangmingensis | 4.51 | 2.17 | 1.50E-06 | 5.82 |
| A0A380CHW3 | Uncharacterized protein conserved in bacteria | *Sporosarcina pasteurii (Bacillus pasteurii)* | 8.67 | 3.12 | 1.54E-06 | 5.81 |

| | | | | | | |
|---|---|---|---|---|---|---|
| A0A560UVL1 | NCS1 family nucleobase:cation symporter-1 | *Burkholderia sp.* SJZ115 | 3.45 | 1.79 | 1.61E-06 | 5.79 |
| R6CXZ9 | Magnesium chelatase 67 kDa subunit ChlI | *Clostridium sp*. CAG:242 | 6.01 | 2.59 | 3.76E-06 | 5.43 |
| A0A1V6WNV2 | Epimerase domain-containing protein | *Penicillium nalgiovense* | 3.65 | 1.87 | 4.42E-06 | 5.35 |
| A0A653AVX6 | RNA polymerase-associated protein RapA (EC 3.6.4.-) | *Clostridium neonatale* | 3.64 | 1.86 | 6.43E-06 | 5.19 |
| A0A024C880 | UvrABC system protein C (Protein UvrC) | *Helicobacter pylori* (*Campylobacter pylori*) | 7.43 | 2.89 | 1.09E-05 | 4.96 |
| A0A0Q8LUI8 | FAS1 domain-containing protein | *Microbacterium* sp. Root180 | 2.74 | 1.45 | 1.13E-05 | 4.95 |
| A0A328KIF3 | Mtd_N domain-containing protein | *Dolosigranulum pigrum* | 2.80 | 1.49 | 1.27E-05 | 4.90 |
| A0A1H6SCG9 | Tfp pilus assembly protein PilF | *Pseudomonas sp.* NFR16 | 5.88 | 2.56 | 1.52E-05 | 4.82 |
| A0A5N6TPT2 | Serine/threonine-protein kinase Tel1 (EC 2.7.11.1) | *Aspergillus avenaceus* | 3.84 | 1.94 | 1.94E-05 | 4.71 |
| T5JHK1 | Phosphoesterase | *Lactiplantibacillus plantarum* EGD-AQ4 | 3.39 | 1.76 | 2.33E-05 | 4.63 |
| A0A4Y8C8N0 | DUF2603 domain-containing protein | *Campylobacter* sp. CH185 | 2.05 | 1.03 | 3.84E-05 | 4.42 |
| A0A1R1QL21 | GNAT family N-acetyltransferase | *Bacillus swezeyi* | 3.21 | 1.68 | 4.11E-05 | 4.39 |
| A0A146FR87 | RTM1-like protein | *Aspergillus kawachii* | 2.40 | 1.26 | 5.68E-05 | 4.25 |
| A0A6G6HNR2 | - | - | 0.09 | -3.46 | 5.74E-05 | 4.24 |
| E6K4I4 | Antioxidant, AhpC/TSA family (EC 1.11.1.15) | *Prevotella buccae* ATCC 33574 | 2.49 | 1.32 | 5.94E-05 | 4.23 |

| | | | | | | |
|---|---|---|---|---|---|---|
| A0A7G1JC81 | - | - | 2.83 | 1.50 | 6.92E-05 | 4.16 |
| A0A3M0LX63 | Zinc ribbon domain-containing protein | *Lactobacillus sp.* ESL0246 | 3.64 | 1.86 | 7.62E-05 | 4.12 |
| A0A2S3WW57 | NADPH quinone reductase MdaB | *Pseudomonas putida* (*Arthrobacter siderocapsulatus*) | 3.39 | 1.76 | 0.000103 | 3.99 |
| A0A4Q4RJE8 | Putative peptide transporter ptr2 | Alternaria arborescens | 3.96 | 1.99 | 0.00012 | 3.92 |
| A0A420VMA9 | Methyl-accepting chemotaxis protein | *Campylobacter* sp. P255 | 2.45 | 1.30 | 0.000124 | 3.91 |
| A0A5E7E2V0 | - | - | 2.98 | 1.57 | 0.000133 | 3.88 |
| A0A2B0MYX4 | DUF3131 domain-containing protein | *Bacillus cereus* | 2.03 | 1.02 | 0.000259 | 3.59 |
| A0A1L9NA58 | Mus7/MMS22 family-domain-containing protein | Aspergillus tubingensis (strain CBS 134.48) | 2.74 | 1.45 | 0.000386 | 3.41 |
| A0A4P9DWZ3 | - | - | 2.22 | 1.15 | 0.000516 | 3.29 |
| A0A2I1MBR3 | Rhodanese domain-containing protein | *Anaerococcus octavius* | 4.08 | 2.03 | 0.00179 | 2.75 |
| I1CN50 | N-acetyltransferase domain-containing protein | *Rhizopus delemar* (*Rhizopus arrhizus var. delemar*) | 2.53 | 1.34 | 0.002608 | 2.58 |
| A0A1T4MIG4 | NAD kinase (EC 2.7.1.23) (ATP-dependent NAD kinase) | Eubacterium ruminantium | 2.84 | 1.50 | 0.004736 | 2.32 |
| A0A150FPS1 | Asp23/Gls24 family envelope stress response protein | [*Clostridium*] *paradoxum* JW-YL-7 = DSM 7308 | 2.08 | 1.06 | 0.006454 | 2.19 |
| A0A1C5QLC9 | DNA primase (EC 2.7.7.101) | uncultured *Clostridium* sp | 0.31 | -1.71 | 0.006713 | 2.17 |
| A0A6G6HN10 | - | - | 2.55 | 1.35 | 0.006797 | 2.17 |

| | | | | | | |
|---|---|---|---|---|---|---|
| A0A2D0ABQ9 | Immunoglobulin lambda-1 light chain-like | *Microbacterium* sp. AISO3 | 6.82 | 2.77 | 0.010176 | 1.99 |
| A8MKW3 | DUF364 domain-containing protein | *Alkaliphilus oremlandii* (strain OhILAs) (*Clostridium oremlandii* (strain OhILAs)) | 6.21 | 2.63 | 0.010784 | 1.97 |
| A0A244ETL9 | Short-chain dehydrogenase | *Pseudomonas syringae* | 0.01 | -6.12 | 0.013377 | 1.87 |
| A0A2X2PC29 | GCN5-related N-acetyltransferase (EC 2.3.1.-) | *Burkholderia cepacia* (*Pseudomonas cepacia*) | 2.22 | 1.15 | 0.015052 | 1.82 |
| D6GT28 | Cation-transporting ATPase, E1-E2 family | *Filifactor alocis* (strain ATCC 35896 / D40 B5) (*Fusobacterium alocis*) | 2.25 | 1.17 | 0.023691 | 1.63 |
| A0A660LVN9 | Nicotinate phosphoribosyltransferase (EC 6.3.4.21) | *Campylobacter sp* | 0.17 | -2.54 | 0.027219 | 1.57 |
| A0A2D3LJD2 | Peptidase | *Prevotella intermedia* | 0.19 | -2.36 | 0.03052 | 1.52 |
| A0A6C0TPP5 | DUF1311 domain-containing protein | *Pseudomonas sp*. OIL-1 | 2.78 | 1.47 | 0.045154 | 1.35 |

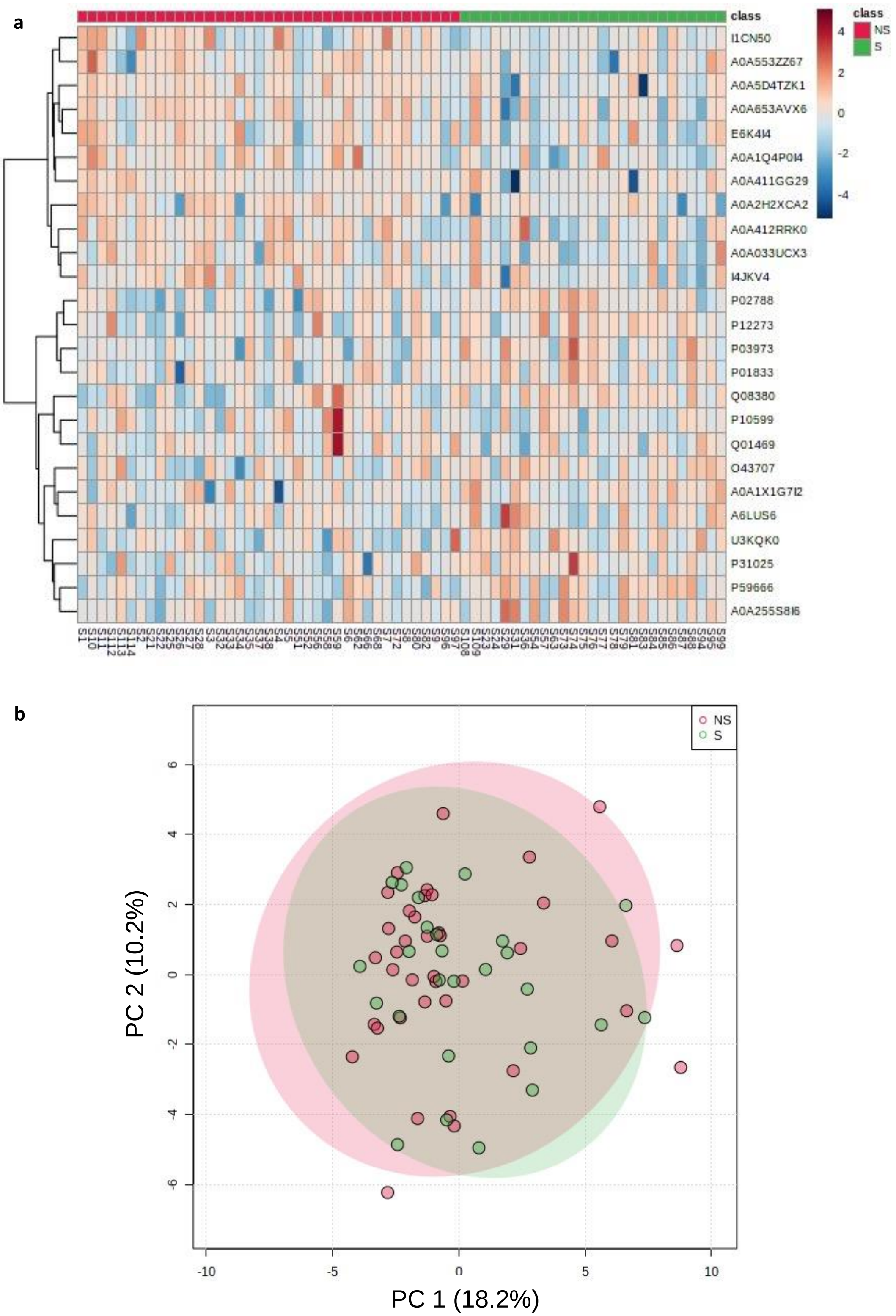


**Figure S4:** Metaproteomic analysis of severe (S) and non-severe (NS) COVID-19 patients from second wave of COVID-19 in India (2021)**: (a)** Hierarchical clustering heatmap and **(b)** PCA plot for severe and non-severe group comparison.

**Section 8. PARALLEL REACTION MONITORING OF MICROBIAL PEPTIDES:**

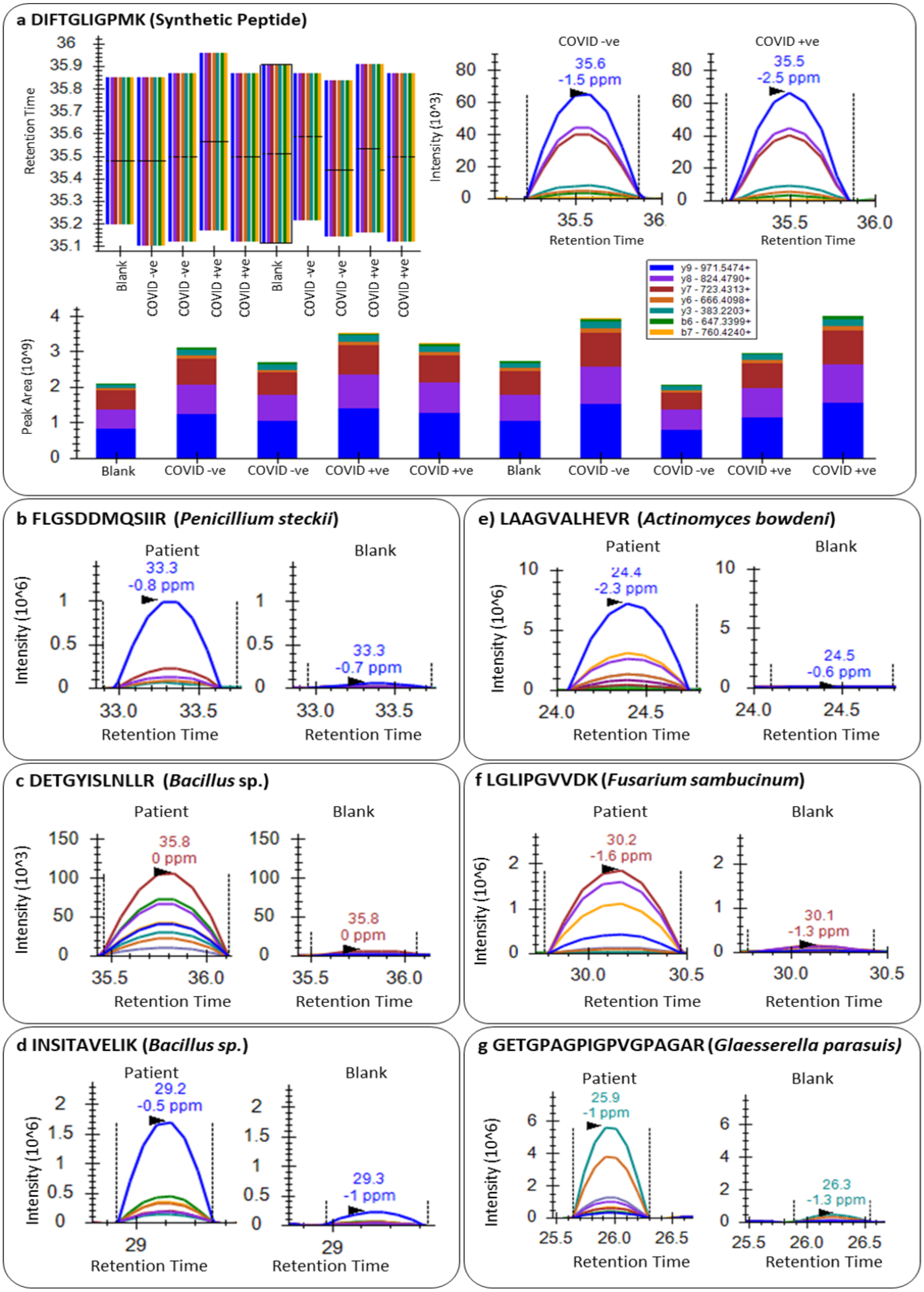

**Figure S5: Detection of microbial peptides in nasopharyngeal swab samples by parallel reaction monitoring (PRM):** (a) retention time (top left), representative peaks (top right), and peak area of the synthetic peptide "DIFTGLIPMK" in the samples. (b), (c), (d), (e), (f), and (g) show peak intensities for peptides FLGSDDMQSIIR, DETGYISLNLLR, INSITAVELIK, LAAGVALHEVR, LGLIPGVVDK, and GETGPAGPIGPVGPAGAR respectively in patient sample vs blank.
